# Human subsystems of medial temporal lobes extend locally to amygdala nuclei and globally to an allostatic-interoceptive system

**DOI:** 10.1101/659854

**Authors:** Adriana L. Ruiz-Rizzo, Florian Beissner, Kathrin Finke, Hermann J. Müller, Claus Zimmer, Lorenzo Pasquini, Christian Sorg

## Abstract

In mammals, the hippocampus, entorhinal, perirhinal, and parahippocampal cortices (i.e., core regions of the human medial temporal lobes, MTL) are locally interlaced with the adjacent amygdala nuclei at the structural and functional levels. At the global brain level, the human MTL has been described as part of the default mode network whereas amygdala nuclei as parts of the salience network, with both networks forming collectively a large-scale brain system supporting allostatic-interoceptive functions. We hypothesized (i) that intrinsic functional connectivity of slow activity fluctuations would reveal human MTL subsystems locally extending to the amygdala; and (ii) that these extended local subsystems would be globally embedded in large-scale brain systems supporting allostatic-interoceptive functions. From the resting-state fMRI data of three independent samples of cognitively healthy adults (one main and two replication samples: Ns = 101, 61, and 29, respectively), we analyzed the functional connectivity of fluctuating ongoing BOLD-activity within and outside the amygdala-MTL in a data-driven way using masked independent component and dual-regression analyses. We found that at the local level MTL subsystems extend to the amygdala and are functionally organized along the longitudinal amygdala-MTL axis. These subsystems were characterized by a consistent involvement of amygdala, hippocampus, and entorhinal cortex, but a variable participation of perirhinal and parahippocampal regions. At the global level, amygdala-MTL subsystems selectively connected to salience, thalamic-brainstem, and default mode networks – the major cortical and subcortical parts of the allostatic-interoceptive system. These results provide evidence for integrated amygdala-MTL subsystems in humans, which are embedded within a larger allostatic-interoceptive system.

## 1 Introduction

The human medial temporal lobes (MTL) include the hippocampal, entorhinal, perirhinal, and parahippocampal cortices (Squire et al., 2004). An extensive body of research has focused on these core MTL regions’ internal (i.e., local) and external (i.e., global) connectivity and how they underpin distinct cognitive functions such as declarative memory or spatial navigation (e.g., Squire et al., 2004; Buzsaki and Moser, 2013; Strange et al., 2014). Specifically, distinct subsystems largely invariable across mammals have been identified within the core MTL (Strange et al., 2014). However, based on anatomy, structural connectivity, and functional interactions, these subsystems consistently appear to extend beyond the core MTL, for example, to the amygdala nuclei. The current study used intrinsic connectivity of slowly fluctuating ongoing activity, to examine in humans for the presence of a ‘local extension’ of core MTL subsystems to the amygdala, and to explore the corresponding ‘global extension’ of the (extended) core MTL-amygdala subsystems to the rest of the brain.

At the local level, in rodents and non-human primates, core MTL regions are highly interrelated with adjacent amygdala nuclei regarding anatomical proximity (Van Hoesen, 1995; Murray and Wise, 2004), structural connectivity (Pitkanen et al., 2000; Petrovich et al., 2001; Kemppainen et al., 2002), and functional interactions (Davis, 1992; Phelps, 2004; Gross and Canteras, 2012). Core MTL regions and the amygdala are anatomically adjacent in mammals (Insausti, 1993; McDonald, 1998), and, in particular in primates, are considered integral components of the MTL as a whole (Van Hoesen, 1995; Amunts et al., 2005). Concerning structural connectivity, dense reciprocal connections exist between the amygdala and core MTL regions (Saunders and Rosene, 1988). In the rat, for example, lateral, basal, and posterior cortical nuclei of the amygdala provide segregated, parallel, and point-to-point organized inputs to parahippocampal, entorhinal, and hippocampal cortices (Pitkanen et al., 2000; Petrovich et al., 2001; Kemppainen et al., 2002), with these topographically organized connections being highly conserved across species (Amaral and Insausti, 1992; Sah et al., 2003). Moreover, functional interaction takes place in a wide variety of functional domains, such as in fear-related unconditioned responses (Davis, 1992; Gross and Canteras, 2012), and emotional (Davis et al., 1994; Gross and Canteras, 2012; Tovote et al., 2015) or episodic memory (Kemppainen et al., 2002; Dolcos et al., 2004).

In humans, intrinsic connectivity has been applied to study local subsystems across either amygdala nuclei or core MTL regions (e.g., Libby et al., 2012; Oler et al., 2012; Maass et al., 2015). Intrinsic connectivity is defined as functional connectivity (iFC) of ongoing slowly fluctuating brain activity (below 0.1 Hz), which is typically measured by correlated blood oxygenation levels of resting-state functional MRI (rs-fMRI) (Fox and Raichle, 2007; Smith et al., 2009). Regarding the amygdala, significant iFC has been found between the centromedial nuclei and the bed nucleus of the stria terminalis in both humans and macaques (Oler et al., 2012). Concerning the core regions of the MTL, an anterior-posterior iFC gradient has been found between the anterior-lateral entorhinal and perirhinal cortices and the proximal subiculum, as well as between the posterior-medial entorhinal and parahippocampal cortices and the distal subiculum (Maass et al., 2015). A comparable gradient-like local organization has also been reported for the hippocampus (e.g., Blessing et al., 2016).

At the global level in humans, iFC has also been applied to study intrinsic connectivity of amygdala nuclei or the core MTL to the rest of the brain (Kahn et al., 2008; Etkin et al., 2009; Roy et al., 2009; Ranganath and Ritchey, 2012; Navarro Schroder et al., 2015; Qin et al., 2015; Wang et al., 2016). For example, whereas basolateral amygdala nuclei link preferentially with the temporal and medial frontal cortices (the ‘fronto-temporal’ amygdala network; Etkin et al., 2009; Roy et al., 2009; Fox et al., 2015), the centromedial nuclei are functionally connected to the midbrain, thalamus, and cerebellum (Etkin et al., 2009). Similarly, the anterior-lateral entorhinal cortices connect with the medial-prefrontal and orbitofrontal cortices, whereas the posterior-medial entorhinal cortices are preferentially connected with posterior parietal areas (Navarro Schroder et al., 2015). Comparable distinct global iFC patterns have also been described for the hippocampus (Kahn et al., 2008; Robinson et al., 2015). At the large-scale system level, the amygdala and the core MTL have been respectively described as part of the salience network – which also includes the insula, the anterior cingulate, and the hypothalamus (e.g., Seeley et al., 2007) – and the default mode network – comprising the precuneus, posterior cingulate, angular gyrus, and medial prefrontal cortex (e.g., Buckner et al., 2008). Recently, both networks have been suggested to be part of a large-scale brain system that appears to support distinct functional domains (i.e., emotion, memory, and social cognition) and is thought to link the control of homeostatic body-focused (i.e., interoceptive) processes with the control of interactions with the environment (i.e., allostatic; Barrett and Simmons, 2015; Kleckner et al., 2017). Beyond the salience and default mode networks, this allostatic-interoceptive system includes key subcortical regions such as parts of striatum, pallidum, thalamus, hypothalamus, and upper brainstem.

On this background, the current study examined for a local and a global extension of the core MTL to the amygdala (i.e., ‘A-MTL’) in humans, as defined by iFC. At the local level, based on structural connectivity evidence of A-MTL subsystems in rodents and primates, we expected analogous subsystems characterized by iFC consistently spanning core regions of the MTL *and* the amygdala. At the global level, based on the respective iFC patterns of the core MTL and the amygdala with networks contributing to the allostatic-interoceptive system, we expected that A-MTL subsystems collectively extend to the allostatic-interoceptive system through iFC. To test these hypotheses, we assessed 101 young healthy participants and analyzed the intrinsic connectivity of fluctuating ongoing rs-fMRI activity within and outside the A-MTL in a data-driven way using masked independent component analysis (Beissner et al., 2014; Blessing et al., 2016) and dual regression (Beckmann et al., 2009; Filippini et al., 2009). Specifically, our mask of independent component analysis was centered on the A-MTL to reveal A-MTL subsystems on the one hand, and their global extension on the other. To control the reliability of our findings, we replicated our approach in two further independent samples.

## 2 Materials and Methods

### 2.1 Participants

One hundred and one healthy young participants (age: 26.7 ± 0.7 years, range: 25-27, 41 females) underwent resting-state fMRI at the Department of Neuroradiology, Klinikum rechts der Isar, Munich, Germany. The local ethics committee of the Klinikum rechts der Isar approved the study, and all participants gave written informed consent for their participation. Study exclusion criteria were current or past neurological and psychiatric disorders, as well as severe systemic diseases or neurotropic medication. The rs-fMRI data of two additional samples of cognitively normal adults participating in other studies (one from our group, and one from a public data base) were used for replication analyses (Replication sample 1: N = 61, age: 36.4 ± 13.6 years, range: 18-65, 18 females. Replication sample 2: N = 29, age: 26.0 ± 4.1 years, range: 18-35, 14 females; see Supplementary Material for more details).

### 2.2 MRI Data Acquisition

MRI data acquisition of the main sample was performed on a Philips Achieva 3T TX system (Netherlands), using an 8-channel SENSE head coil. Functional MRI T2*-weighted data were collected for 10 min 52 s while participants were resting with eyes closed, and after being instructed not to fall asleep. We verified that subjects stayed awake by interrogating via intercom immediately after the resting-state fMRI scanning run. Two hundred and fifty volumes of blood oxygenation level dependent (BOLD) rs-fMRI signal per individual were acquired using a gradient-echo echo planar imaging (GRE-EPI) sequence: Repetition time, TR = 2,608 ms; echo time, TE = 35 ms; phase encoding direction: anterior–posterior; flip angle = 90°; field of view, FOV = 230 mm; matrix size = 64 × 64, 41 interleaved slices, and no interslice gap; reconstructed voxel size = 3.59 mm isotropic. Subsequently, a high-resolution T1-weighted image was acquired using a 3D-MPRAGE sequence with the following parameters: TR = 7.7 ms; TE = 3.9 ms; inversion time, TI = 1,300 ms; flip angle = 15°; 180 sagittal slices, reconstruction matrix: 256 × 256; reconstructed voxel size 1 mm isotropic.

### 2.3 Data Preprocessing

Functional MRI data were preprocessed using the Data Processing Assistant for Resting-State fMRI toolbox (DPARSF; Chao-Gan and Yu-Feng, 2010) and SPM12 (http://www.fil.ion.ucl.ac.uk/spm). After discarding the first 5 rs-fMRI volumes to avoid magnetization effects, functional volumes were realigned to correct for head motion. Each participant’s T1-weighted structural image was segmented into gray matter, white matter, and cerebrospinal fluid using the tissue classification algorithm implemented in SPM, which is based on prior probabilities of voxels belonging to each tissue type (obtained from scans of 152 healthy young subjects provided by the Montreal Neurological Institute, MNI). Each participant’s rs-fMRI volumes were coregistered to their high-resolution structural T1 image by using boundary-based registration, and then transformed to MNI space at 2 × 2 × 2-mm^3^ resolution using nonlinear registration derived from T1 image normalization and then spatially smoothed using a 5 mm full-width-at-half-maximum (FWHM) Gaussian kernel. To control for movement artifacts, we used as criterion peak-to-peak motion below 1 mm or 1 degree in any direction. Based on this criterion, all subjects were included in further analyses. To control for nuisance covariates, we extracted the mean time series for white matter (WM) and cerebrospinal fluid (CSF) from the fMRI data. Each individual’s segmented high-resolution structural MRI was used to calculate WM and CSF specific mean time series, with tissue type probability of 0.8, by averaging across all voxels within the tissue masks. WM, CSF, and global signals, and six head motion parameters (three translations, three rotations) for each subject were regressed out from the fMRI data.

### 2.4 Data Analysis

We analyzed the preprocessed rs-fMRI data by employing masked independent component analysis (mICA) combined with dual-regression analysis (Beissner et al., 2014; Blessing et al., 2016). Masked ICA localized iFC-based sources within a mask (i.e., the local A-MTL subsystems), whereas whole-brain dual regression identified corresponding global-iFC patterns related to the local sources. A mask of the A-MTL was built by combining the bilateral masks of the amygdala, hippocampus, and entorhinal, perirhinal, and posterior parahippocampal cortices, derived from the Harvard-Oxford cortical and subcortical probabilistic structural atlases and the Jülich histological atlas, using Fslview (http://fsl.fmrib.ox.ac.uk/fsl/fslview/). These masks were added up using the Imcalc toolbox of SPM (http://www.fil.ion.ucl.ac.uk/spm/), binarized at a threshold probability of 0.5, and resampled to the size of our functional data. Next, the preprocessed fMRI data were temporally concatenated and analyzed by probabilistic ICA (Beckmann and Smith, 2004) restricted to the A-MTL mask, using the FSL *melodic* command (http://www.fmrib.ox.ac.uk/fsl/). First, these data were normalized for voxel-wise mean and variance, and then reduced into a 20-dimensional subspace by probabilistic principal component analysis. A dimensionality of 20 was chosen based on a series of control analyses, which are described in detail below. Subsequently, data were decomposed into time courses and spatial maps by optimizing for non-Gaussian spatial distributions using a fixed-point iteration technique (Hyvarinen, 1999). The resulting group-level component maps are divided by the standard deviation of the residual noise and thresholded by fitting a mixture model to the histogram of intensity (Beckmann and Smith, 2004).

### 2.5 Dual regression

To assess both local- and global-iFC of A-MTL subsystems, dual-regression analyses were conducted (Beckmann et al., 2009; Filippini et al., 2009; Smith et al., 2014; Nickerson et al., 2017). Dual regression is a multivariate approach that allows the estimation of an *individual* version of the group-level spatial maps. Dual regression works in two steps. In the first step, the set of spatial independent components derived by group-level mICA is regressed on the individual participant’s 4D dataset in a multiple regression. This results in a set of participant-specific time courses, one per group-level spatial map. In the second step, those time courses are regressed in a second multiple regression, on the same 4D dataset, resulting in participant-specific spatial maps, one per group-level spatial map. Participant-specific spatial maps were further analyzed in two ways: (i) maps were restricted to the A-MTL mask and used to estimate local-iFC patterns and iFC peaks of A-MTL; (ii) a whole-brain mask was used to estimate the global-iFC patterns that correspond to those local-iFC patterns. Next, the statistical significance of both restricted and unrestricted maps was assessed in a two-sided one-sample t-test using FSL’s *randomise* permutation-testing tool, resulting in p-value maps for each component involved in the analysis. Specifically, the results were based on 500 permutations and a p-value of 0.05, corrected for multiple comparisons by threshold-free cluster enhancement (Smith and Nichols, 2009).

### 2.6 Separation of neural from non-neural local-iFC patterns

To separate local-iFC patterns of ‘neural’ and ‘non-neural’ origin, we analyzed their associated global-iFC patterns to compute, for each one of them, the percentage of voxels that lay on gray matter (GM), WM, and CSF. We used the tissue probability maps of the three tissue types of interest (downloaded from: https://www.fil.ion.ucl.ac.uk/spm/toolbox/TPM/) (Blaiotta et al., 2018). The thresholded (to a probability of 0.9) maps were multiplied with the thresholded (voxels greater than 0.95) and binarized p-value maps of our mICA independent components (following Beissner et al., 2014). From this multiplication, we obtained, for each global-iFC pattern, the number of voxels present in each tissue type (i.e., GM, WM, and CSF). Finally, we calculated, for each global-iFC pattern, the percentage of voxels in CSF with respect to the total number of voxels (i.e., in the three tissue types; based on the approach of Beissner et al., 2014). Based on the mean percentage of voxels in CSF (from all global-iFC patterns), we tagged as ‘non-neural’ those global-iFC patterns with a percentage at or above the mean and as ‘neural’ those below the mean (see Table S1). Our results are thus based only on those iFC patterns classified as ‘neural.’

### 2.7 Anatomical characterization of local-iFC patterns

After excluding non-neural local-iFC patterns, we characterized the remaining maps anatomically. Specifically, the iFC peaks of these maps were identified using the FSL tool *fslstats*. Peaks were classified as anterior (y = 4 to −18 mm), middle (y = −19 to −31 mm), or posterior (y = −32 to −42 mm) [following the classification of the longitudinal axis of whole MTL by Kivisaari et al. (2013)]. The extent of iFC (i.e., involvement of each A-MTL structure) was also examined for each map and slice by slice in the coronal plane.

### 2.8 Local-iFC patterns and associated large-scale brain networks

Based on previous findings relating local-iFC subsystems of core MTL regions and amygdala to known large-scale brain networks (Kahn et al., 2008; Etkin et al., 2009; Roy et al., 2009; Ranganath and Ritchey, 2012; Navarro Schroder et al., 2015; Qin et al., 2015; Wang et al., 2016), we expected that the global-iFC of A-MTL subsystems would also be embedded in the functional architecture of large-scale brain networks (Allen et al., 2011; Yeo et al., 2011; Raichle, 2015) – particularly those constituting the allostatic-interoceptive system, i.e., the default mode, salience, and brainstem-thalamus networks. Thus, we compared the global-iFC patterns of the ‘neural’ local-iFC patterns with templates of large-scale brain networks derived from a study that estimated these networks from rs-fMRI data of about 600 healthy participants (Allen et al., 2011). We calculated spatial cross-correlation coefficients between those templates and our global-iFC patterns using the *fslcc* command in FSL (Jenkinson et al., 2012). The global-iFC pattern with the highest cross-correlation coefficient was chosen as the pattern best matching a particular known large-scale brain network.

### 2.9 Control analyses: number of ICA dimensions and replication

Since ICA results depend heavily on the number of dimensions used to decompose the data, we searched for the optimal dimensionality in our specific A-MTL mICA approach. Specifically, we performed a series of control analyses of three ICA dimensionalities, 10, 20, and 30. We defined the ‘optimal dimensionality’ based on the neural global-iFC patterns, whereby ‘optimal’ meant that the global-iFC patterns were separated into a maximum number without generating qualitatively new iFC-patterns, but also without generating redundant iFC patterns.

Second, to control both the reliability of our findings and the impact of smaller sample size, we replicated our approach in two further independent samples. We used control samples with lower sample sizes (one medium sized and one small sized) compared to that used in the present study (replication sample 1: N = 61; replication sample 2: N = 29). Moreover, the size of one of the samples approached more closely the sample size typically used in patient studies (i.e., ∼ 30). Both replication datasets have been previously described and used for other analyses (see Supplementary Material for details). Using as well the optimal mICA dimensionality and selection of the local-iFC patterns via global-iFC patterns, we then tested their respective cross-correlations with the original results, as described in more detail in the Supplementary Material.

## 3 Results

### 3.1 Neural iFC patterns, optimal number of dimensions for mICA, and naming of iFC patterns

Figure 1 shows twelve local-iFC patterns and their corresponding global-iFC patterns identified as of ‘neural’ origin for the mICA with dimensionality of 20 (see below). Figure S1 shows the remaining 8 local- and global-iFC patterns rated as ‘non-neural’ (see Table S1 for percentage of voxels in CSF in these patterns).

**Figure 1.**
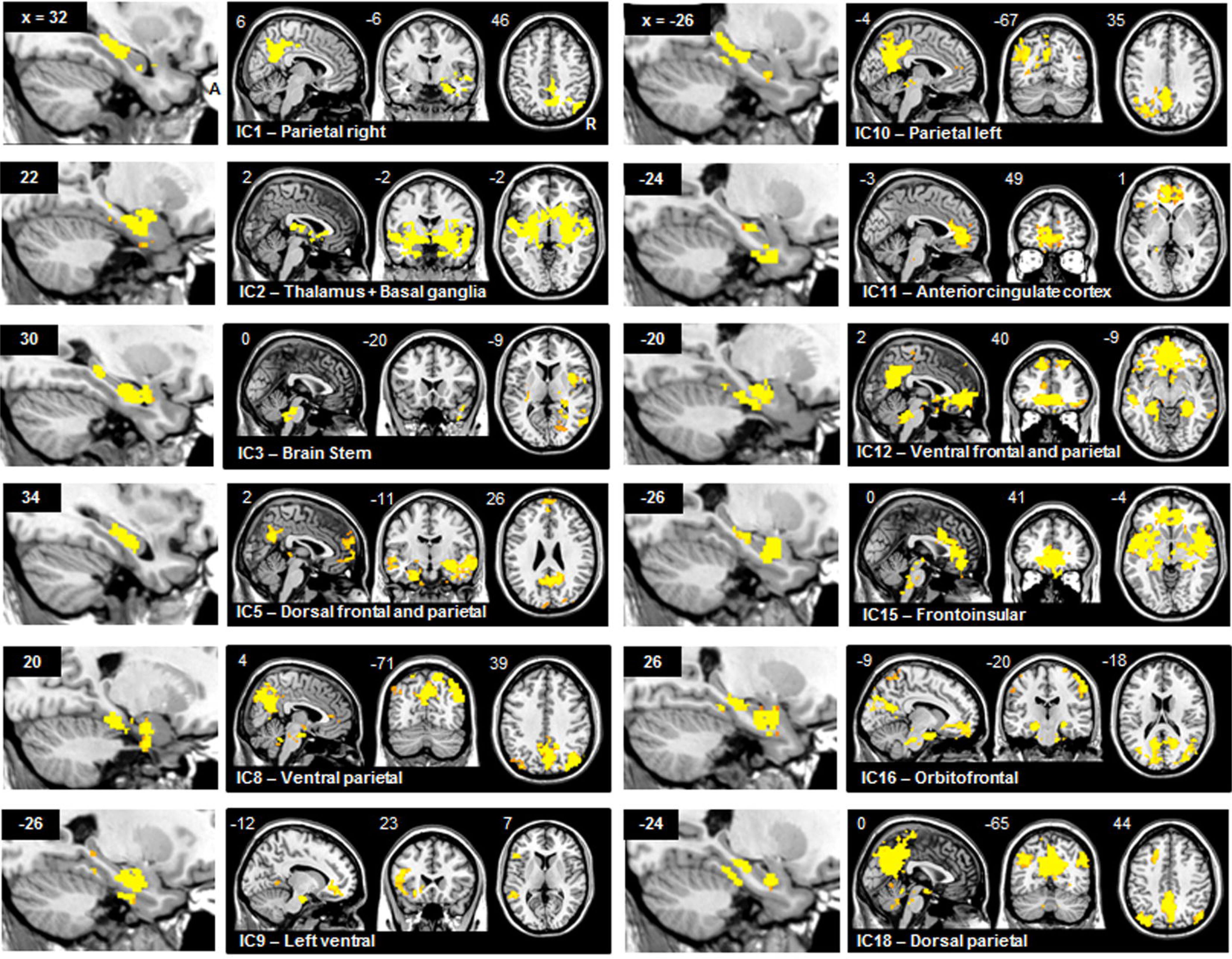
A-MTL local- and global-iFC patterns. A-MTL iFC patterns identified as of neural origin and obtained from the masked independent component analysis (mICA) with 20 dimensions and subsequent whole-brain dual regression. In yellow, significant voxels of the local- and global-iFC patterns (p-values; p < 0.05 FWE corrected for multiple comparisons). MNI coordinates in mm (x, y, z) are shown on the left upper corner of each image. Right is shown on the right of axial and coronal images; anterior is shown on the right of sagittal images. A: anterior; IC: independent component; R: right.

We selected a dimensionality of 20 as optimal for the mICA because neural global-iFC patterns tended to merge in the 10-dimension mICA, but to split (without providing new information) or repeat in the 30-dimension mICA (Table S2). For example, a global-iFC pattern involving frontal and parietal cortices in 10-mICA (Figure S2,) could be split into three global-iFC patterns in 20-mICA, with two involving lateral parietal cortices and one involving medial frontal and parietal cortices. These three global-iFC patterns could, in turn, also appear in the 30-mICA but with additional redundant or uninformative iFC patterns (see also Table S2 and Table S3).

We named A-MTL global-iFC patterns according to the regions they included, as a way to identify them without implying any a priori function (Figure 1 and last column of Table 1). These A-MTL global-iFC patterns comprised regions of and corresponded well to large-scale brain networks reported in the literature (Table 2). For example, one of our global-iFC patterns corresponded to the “salience network” of Allen et al. (2011) (IC15 – Frontoinsular), one to “basal ganglia” (IC2 – Thalamus + basal ganglia), and at least seven of our global-iFC patterns to the “DMN posterior lateral.”

**Table 1.** Local-iFC patterns organized along the longitudinal axis based on the peak iFC.

| Coordinates<br>(x, y, z) of<br>the peak<br>voxel in MNI<br>space | Local-iFC<br>pattern | Brain structures of local-iFC A-MTL patterns |  |  | Brain structures of global-iFC<br>patterns | Global-iFC<br>pattern label |
| --- | --- | --- | --- | --- | --- | --- |
|  |  | Anterior | Central | Posterior |  |  |
| -26, -4, -16 | Anterior AM | ○ AM<br>○ HC<br>○ EC | ○ HC<br>○ EC |  | Anterior cingulate cortex, lateral and medial orbitofrontal cortex, insular cortex, putamen, temporal pole, and supramarginal gyrus, ventral striatum, ventral pallidum, cerebellum (vermis), and periaqueductal gray | Frontoinsular<br>(IC15) |
| 26, -6, -24 | Anterior HC – 1 | ○ AM<br>○ HC<br>○ EC<br>○ PRC | ○ HC<br>○ PHC | ○ PHC | Postcentral gyrus, precuneus, occipital cortex, fusiform cortex, insula, bilateral orbitofrontal cortex, temporal pole | Orbitofrontal<br>(IC16) |
| -24, -8, -36 | Anterior PRC | ○ AM<br>○ HC<br>○ EC<br>○ PRC | ○ HC<br>○ PHC | ○ PHC | Medial orbitofrontal cortex, fronto-insular cortex, temporal lobe, ventral striatum, and putamen | Anterior<br>cingulate cortex<br>(IC11)* |
| 30, -12, -20 | Anterior HC – 2a | ○ AM<br>○ HC<br>○ EC<br>○ PRC | ○ HC<br>○ EC | ○ HC<br>○ PHC | Brain stem, cerebellum, thalamus, middle temporal gyrus, insula, fusiform cortex, precentral gyrus, inferior lateral occipital cortex | Brain stem<br>(IC3) |
| 22, -12, -18 | Anterior HC – 2b | ○ AM<br>○ HC<br>○ EC<br>○ PRC | ○ HC<br>○ PHC | ○ HC<br>○ PHC | Insular cortex, superior temporal gyrus, nucleus accumbens, caudate nucleus, putamen, and thalamus | Thalamus +<br>basal ganglia<br>(IC2) |
| -26, -14, -18 | Anterior HC – 3 | ○ AM<br>○ HC<br>○ EC<br>○ PRC | ○ HC<br>○ EC<br>○ PRC<br>○ PHC | ○ HC<br>○ PHC | Left orbitofrontal cortex, left inferior frontal gyrus, left superior and middle temporal gyrus, left thalamus, left caudate, left fusiform cortex, left lateral occipital cortex | Left ventral<br>(IC9) |
| -20, -16, -18 | Anterior HC – 4 | ○ AM<br>○ HC<br>○ EC<br>○ PRC | ○ HC<br>○ EC<br>○ PRC<br>○ PHC | ○ HC<br>○ PHC | Medial frontal cortex, ventromedial frontal cortex, subgenus anterior cingulate cortex, dorsomedial frontal cortex, middle temporal gyrus, retrosplenial cortex, precuneus, posterior cingulate cortex, nucleus accumbens, and lower brain stem | Ventral frontal<br>and parietal<br>(IC12) |
| 34, -20, -14 | Central HC – 1 | ○ AM<br>○ HC<br>○ EC<br>○ PRC | ○ HC<br>○ PHC | ○ HC<br>○ PHC | Dorsomedial prefrontal cortex, ventromedial prefrontal cortex, anterior part of superior and middle temporal gyrus, nucleus accumbens, precuneus, posterior cingulate cortex, and occipital cortex | Dorsal frontal<br>and parietal<br>(IC5) |
| 32, -26, -14 | Central HC – 2 | ○ AM<br>○ HC<br>○ EC<br>○ PRC | ○ HC<br>○ EC<br>○ PHC | ○ HC<br>○ PHC | Right inferior parietal lobule and posterior cingulate cortex, bilateral precuneus and temporal lobe | Parietal right<br>(IC1) |
| -26, **-28**, -14 | Central HC – 3 | ○ AM<br>○ HC<br>○ EC | ○ HC<br>○ PHC | ○ HC<br>○ PHC | Left inferior parietal lobule and posterior cingulate cortex, bilateral precuneus and temporal lobe | Parietal left<br>(IC10) |
| 20, **-32**, -14 | Posterior PHC – 1 | ○ AM<br>○ HC<br>○ EC<br>○ PRC | ○ HC<br>○ EC<br>○ PRC<br>○ PHC | ○ HC<br>○ PHC | Middle temporal gyrus, paracingulate gyrus, insula, fusiform cortex, inferior parietal lobule, precuneus, anterior cingulate cortex | Ventral parietal<br>(IC8) |
| -24, **-32**, -18 | Posterior PHC – 2 | ○ AM<br>○ HC<br>○ EC | ○ HC<br>○ PHC | ○ HC<br>○ PHC | Bilateral posterior parietal cortex, precuneus, posterior cingulate cortex, hypothalamus, cerebellum, and brain stem | Dorsal parietal<br>(IC18) |

AM = amygdala; EC = entorhinal cortex; HC = hippocampus; iFC = intrinsic functional connectivity; IC = independent component; PHC = posterior parahippocampal cortex; PRC = perirhinal cortex. \*It resembles the semantic appraisal network described in previous studies (Guo et al., 2013; Zhou and Seeley, 2014).

**Table 2.** Spatial cross-correlation between large-scale brain networks and the current global-iFC patterns

| Global-iFC patterns | Allen et al.'s (2011) networks |
| --- | --- |
| IC15 – Frontoinsular | IC55, “salience,” $r = 0.14$ |
| IC16 – Orbitofrontal | IC53, “DMN posterior lateral,” $r = 0.12$ |
| IC11 – Anterior cingulate cortex | IC25, “DMN anterior medial,” $r = 0.26$ |
| IC3 – Brain stem | IC42, “frontal,” $r = 0.11$ |
| IC2 – Thalamus + basal ganglia | IC21, “basal ganglia,” $r = 0.40$ |
| IC9 – Left ventral | IC67, “visual,” $r = 0.11$ |
| IC12 – Ventral frontal and parietal | IC53, “DMN posterior lateral,” $r = 0.28$ |
| IC5 – Dorsal frontal and parietal | IC53, “DMN posterior lateral,” $r = 0.20$ |
| IC1 – Parietal right | IC53, “DMN posterior lateral,” $r = 0.34$ |
| IC10 – Parietal left | IC53, “DMN posterior lateral,” $r = 0.33$ |
| IC8 – Ventral parietal | IC53, “DMN posterior lateral,” $r = 0.35$ |
| IC18 – Dorsal parietal | IC53, “DMN posterior lateral,” $r = 0.48$ |
| Global-iFC patterns with the highest cross-correlation coefficient with Allen et al.'s networks (2011). |  |

### 3.2 Longitudinal organization of A-MTL local- and global-iFC patterns

We arranged the twelve identified neural A-MTL local-iFC patterns based on the location of the iFC peak along the longitudinal axis (Figure 2A and Table 1). In the *anterior* A-MTL (i.e., y = 4 to −18 mm), we observed one local-iFC pattern with an iFC peak in the amygdala, one in the perirhinal cortex, and five in the hippocampus. Regarding the *middle* A-MTL (i.e., y = −19 to −31 mm), three local-iFC patterns were found with distinct iFC peaks along the central hippocampus. Finally, in the *posterior* A-MTL (i.e., y = −32 to −42 mm), two local-iFC patterns were found with iFC peaks in the posterior parahippocampal cortex. No iFC peak was found in the entorhinal cortex or the posterior hippocampus for any of the ‘neural’ local-iFC patterns.

**Figure 2.**
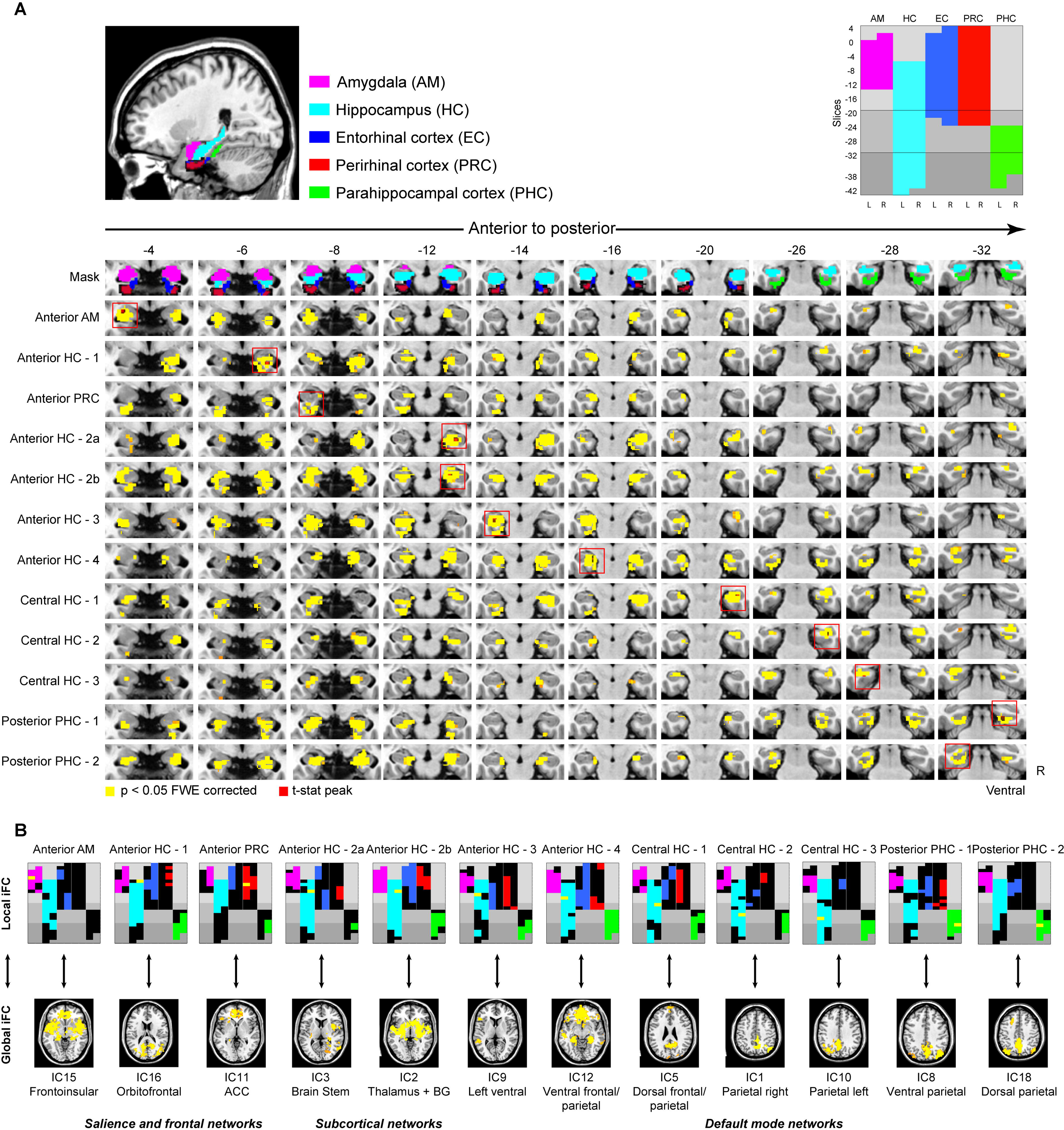
IFC patterns of the A-MTL. **(A)** Anatomical structures of the A-MTL mask used for the mICA and dual regression are color-coded (upper left) and schematically represented, slice-by-slice, along the longitudinal axis (upper right). The local-iFC patterns resulting from the mICA and dual regression are presented below in yellow, organized according to the iFC peak location (surrounded by a red square). The slices were selected to show each local iFC peak and the relative extent of the local-iFC pattern. **(B)** Each local-iFC pattern of **A** is schematically represented following the color coding (upper row) and accompanied by its global-iFC pattern (bottom row). The yellow bar in each schematic local-iFC pattern indicates the location of the iFC peak (shown within the red square in A). The three network labels on the bottom refer to the resemblance of large-scale networks (see Table 2 for details). Slice numbers in A correspond to y coordinates (in mm) in MNI space. Shades of gray in the schemes demarcate anterior, middle, and posterior A-MTL according to Kivisaari et al. (2013). Axial slices of global-iFC patterns are taken from Figure 1. Abbreviations: ACC = Anterior cingulate cortex; BG = basal ganglia; FWE = family wise error; IC = independent component; L = left; R = right.

To visualize and analyze the spatial outline of the twelve identified A-MTL local-iFC patterns in more detail, we mapped each local-iFC pattern onto a longitudinally organized, slice-wise, rectangular schematic of the A-MTL with color-coded ‘boxes’ for amygdala, hippocampus, entorhinal, perirhinal, and parahippocampal cortices (Figure 2B and upper right corner of 2A). Every local-iFC pattern included the different A-MTL structures to different extents (different colors on the black background of the schematics of Figure 2B). However, despite such differential involvement of A-MTL structures in the local-iFC patterns, it is clear from the upper row of Figure 2B that three of those structures were always present (although to different degrees) in every local-iFC pattern, namely: the amygdala, the hippocampus, and the entorhinal cortex (see also Figure S3). This result indicates the consistent involvement of these three structures across the A-MTL local-iFC patterns. In contrast, the perirhinal and posterior parahippocampal cortices did not appear in all local-iFC patterns. Specifically, the perirhinal cortex (in red) was not present in the Anterior AM as well as not in the Central HC - 3 and Posterior PHC - 2 local iFC-patterns. Similarly, the posterior parahippocampal cortex did not appear in the Anterior AM local-iFC pattern.

Next, we added the A-MTL global-iFC patterns to the longitudinal outline of local-iFC patterns. We found a longitudinal gradient for *global*-iFC patterns that remarkably corresponds to that of local-iFC peaks (Figure 2B). In more detail, the most anterior (dorsal) global-iFC patterns – which covered frontoinsular, orbitofrontal, and anterior cingulate cortices, and resembled a frontal and the salience networks of Allen et al. (2011) (Table 2) – corresponded to the three most anterior local-iFC patterns with iFC peaks in the amygdala, anterior hippocampus, and perirhinal cortex, respectively. Subcortical (ventral) global-iFC patterns – which covered brainstem, basal ganglia, and thalamus and resembled the basal ganglia network of Allen et al. (Table 2) – corresponded to the next two anterior local-iFC patterns with two iFC peaks in the anterior hippocampus. Finally, the remaining seven global-iFC patterns corresponded to local-iFC patterns with two iFC peaks in the anterior hippocampus, three in the central hippocampus, and two in the posterior parahippocampal cortex. These seven global-iFC patterns covered anterior and posterior cingulate, dorsal and ventral prefrontal, middle temporal, and inferior parietal cortices as well as lower brain stem, hypothalamus, and cerebellum, and resembled subparts of the default mode network (Table 2). Thus, the anatomical organization of A-MTL local-iFC patterns along the longitudinal axis resulted in a similar organization (i.e., anterior-posterior) of A-MTL global-iFC patterns, which match the large-scale salience and default-mode networks as well as subcortical networks.

### 3.3 Reliability of A-MTL local- and global-iFC patterns

Next, we tested the replicability of A-MTL subsystems. We found similar global-iFC patterns based on mICA restricted to the A-MTL mask in two different samples of cognitively normal adult participants (Figures 3 and 4). For the replication sample 1 (N = 61), 10 global-iFC patterns were identified as ‘neural,’ following the same approach used for the original sample. Of these 10, six had highest spatial cross-correlations with the original sample’s global-iFC patterns: three had the highest spatial cross-correlation with each one of the original brain stem, ventral frontal and parietal, and parietal right; one had the highest cross-correlation with both the frontoinsular and thalamus/basal ganglia iFC patterns of the original sample; one had the highest cross-correlation with the original orbitofrontal and three parietal; and one had it with the original anterior cingulate and left ventral global-iFC patterns (mean coefficient of spatial cross-correlation: 0.30 ± 0.09, range: 0.18 – 0.44; see Figure 3 and Table S4 for details). The local-iFC patterns were also spatially well correlated with the local-iFC patterns of the original sample (mean coefficient: 0.49 ± 0.08, range: 0.37 – 0.60; Figure 4 and Table S4 for details). Finally, the iFC peaks were located less than or equal to 6 voxels in the Y-Z plane (i.e., irrespective of the side) in more than half of them (those corresponding to IC15 – Frontoinsular; IC11 – ACC; IC3 – Brain stem; IC1 – Parietal right; IC10 – Parietal left; IC8 – Ventral parietal; and IC18 – Dorsal parietal).

**Figure 3.**
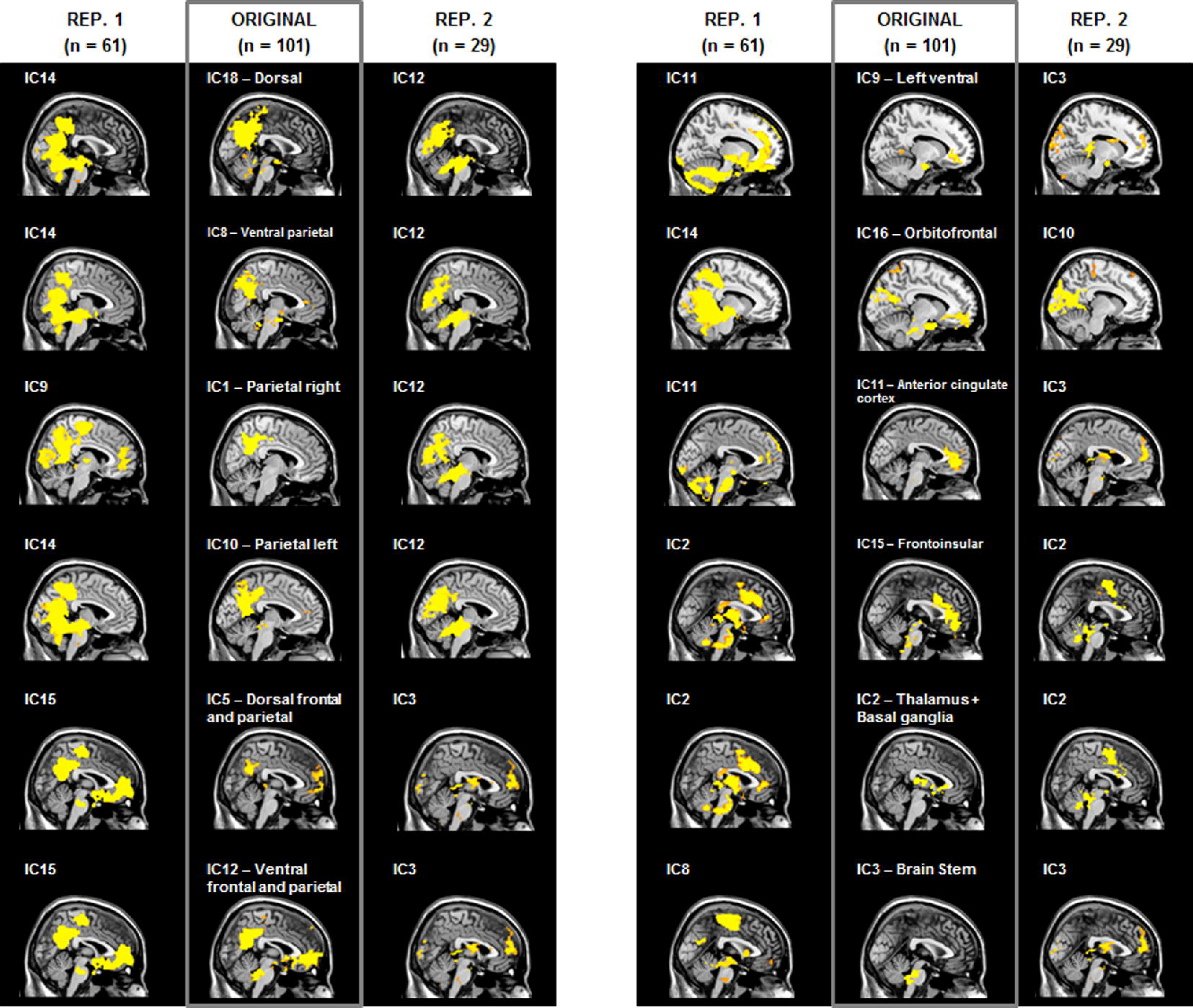
Replication of global-iFC patterns. Global-iFC patterns obtained from masked independent component analysis (mICA) restricted to the medial temporal lobe and amygdala (A-MTL) in three different samples of cognitively normal adults (middle column: original sample, whose results are reported in the main text; first column: images of the replication sample 1, REP. 1, whose details are reported in Table S4; third column: images of the replication sample 2, REP. 2, whose details can be found in Table S5). Sagittal images appear in the same coordinates as those used in the global-iFC patterns of Figure 1.

**Figure 4.**
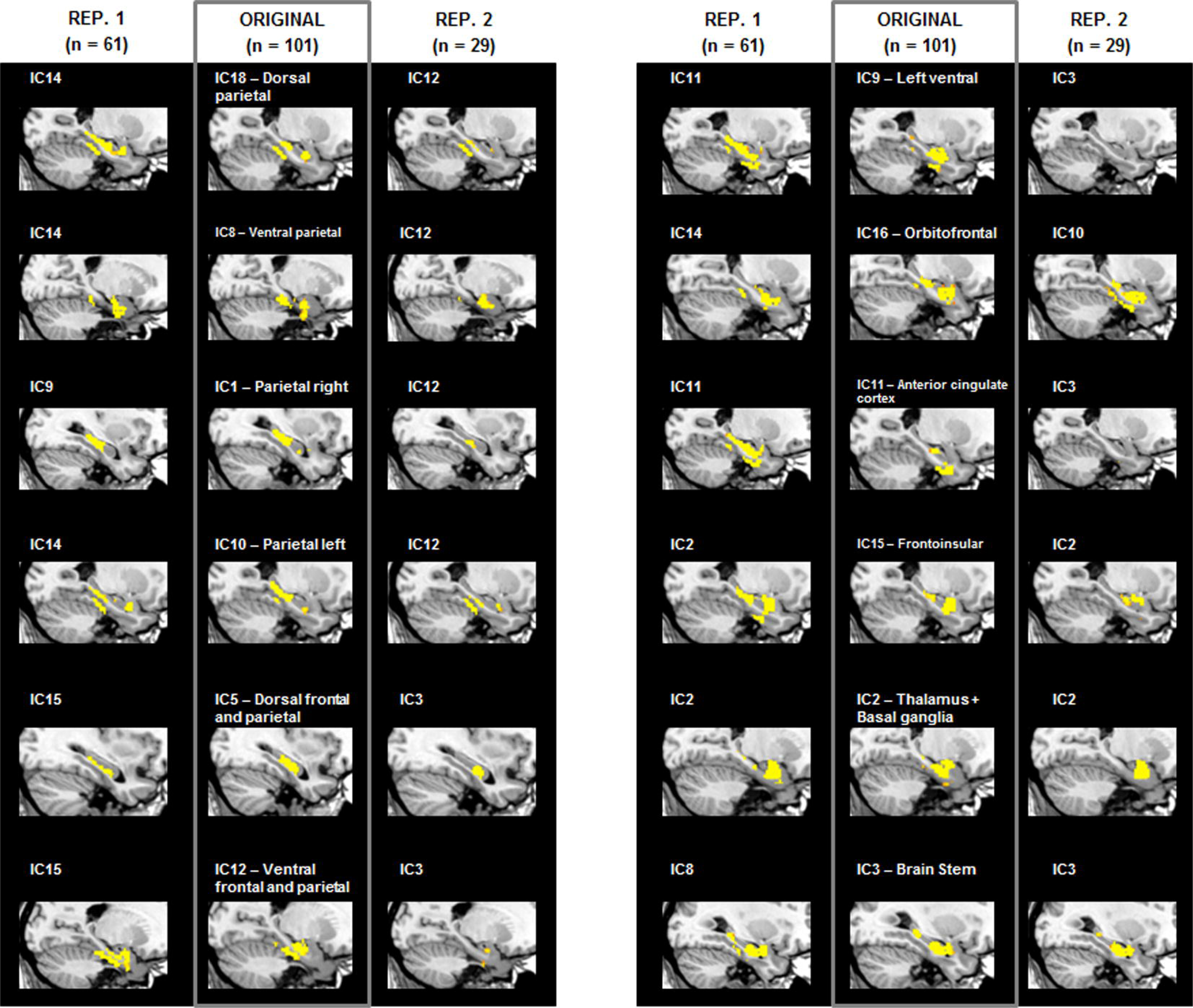
Replication of local-iFC patterns. Local-iFC patterns obtained from masked independent component analysis (mICA) restricted to the medial temporal lobe and amygdala (A-MTL) in three different samples of cognitively normal young adults (middle column: original sample, whose results are reported in the main text; first column: images of the replication sample 1, REP. 1, whose details are reported in Table S4; third column: images of the replication sample 2, REP. 2, whose details can be found in Table S5). Sagittal images appear in the same coordinates as those used in the local-iFC patterns of Figure 1.

For the replication sample 2 (N = 29; Figures 3 and 4), nine independent components were identified as ‘neural.’ Of these 9, four had highest spatial cross-correlations with the original sample’s global-iFC patterns: one global-iFC pattern had the highest spatial cross-correlation with the original orbitofrontal iFC pattern; one had it with the original frontoinsular and thalamus/basal ganglia; one with the original anterior cingulate, brain stem, left ventral, and dorsal frontal and parietal; and one with the five remaining original global-iFC patterns, all of which resembled subparts of the default mode network (mean coefficient of spatial cross-correlation: 0.25 ± 0.08, range: 0.16 – 0.43; see Figure 3 and Table S5 for more details). The corresponding local-iFC patterns were also spatially correlated with the original local-iFC patterns (mean coefficient: 0.43 ± 0.13, range: 0.21 – 0.68; Figure 4 and Table S5 for more details). Finally, as with the first replication sample, more than half of all iFC peaks were located less than or equal to 6 voxels in the Y-Z plane (those corresponding to IC15 – Frontoinsular; IC3 – Brain stem; IC2 – Thalamus + basal ganglia; IC9 – Left ventral; IC1 – Parietal right; IC10 – Parietal left; IC8 – Ventral parietal; and IC18 – Dorsal parietal).

## 4 Discussion

Using a data-driven approach on rs-fMRI data of healthy adults, we identified twelve intrinsic functional connectivity-based subsystems spanning the amygdala and the MTL (A-MTL) with two fundamental properties. First, all subsystems consistently covered parts of the amygdala, the hippocampus, and the entorhinal cortex. Second, subsystems showed a discrete organization along the longitudinal axis of the medial temporal lobes. The distinctive anterior-posterior organization of local connectivity at the A-MTL level is mirrored by a corresponding longitudinal arrangement at the global connectivity level. Specifically, global intrinsic connectivity patterns of A-MTL subsystems are arranged from prefrontal-insular, through subcortical, to posterior cingulate centered patterns. These patterns were remarkably similar to known large-scale brain networks, which are associated with distinct functional domains, proposed to support allostatic-interoceptive functions (Kleckner et al., 2017). These networks were the salience, basal ganglia/thalamus, hypothalamus/brainstem, and default mode networks. Thus, our results provide empirical evidence in humans for distinct A-MTL intrinsic connectivity subsystems with both (i) consistent recruitment of the amygdala, hippocampus, and entorhinal cortex, and (ii) a longitudinal anterior-posterior gradient that corresponds with global-iFC patterns of overlapping insular-cingulate and subcortically centered large-scale brain networks.

### 4.1 A-MTL subsystems derived from A-MTL local- and global-iFC patterns

#### 4.1.1 A-MTL subsystems include the amygdala, the hippocampus, and the entorhinal cortex, and extend along the A-MTL longitudinal axis

One of our primary results is that A-MTL local-iFC patterns consistently span parts of both amygdala and core MTL regions, namely, hippocampus and entorhinal cortex (Figure 2B). Importantly, these local-iFC patterns were independent both from methodological aspects such as mICA dimensionality (Figure S2; Table S2), and sample properties (Figure 4; Tables S4 and S5), thus underscoring the reliability of this finding. Such a combined contribution of the hippocampus and the entorhinal cortex is consistent with the cortical connectivity of the core MTL, with common input into the entorhinal cortex, intra-hippocampal loops, and output via entorhinal cortex (see, e.g., Burwell, 2000; Buzsaki, 2011). Our result is also in line with the previously shown convergence of functional connectivity of both regions, both within and outside the A-MTL, in human neuroimaging studies. For example, anterior parts of both the hippocampus and the entorhinal cortex have been shown to be functionally connected with the lateral temporal cortex (Kahn et al., 2008). Organized functional connectivity has moreover been shown between sub-regions of the entorhinal cortex and the hippocampus on the one hand, and the perirhinal/parahippocampal cortices on the other (Maass et al., 2015). Functional connectivity of the hippocampus, in turn, also extends to the amygdala, in particular to basolateral portions (Roy et al., 2009). Importantly, organized connectivity among the hippocampus, entorhinal cortex, and amygdala is also supported by evidence from structural connectivity, and anatomical, molecular, and functional studies in rodents and primates (Pitkanen et al., 2000; Swanson, 2003; Canto et al., 2008; Strange et al., 2014). For example, reviewed data of connectivity in rodents show substantial reciprocal interconnections between the lateral, basal, and accessory basal nuclei of the amygdala, and the rostral entorhinal cortex and temporal end of the hippocampus (Pitkanen et al., 2000). Functionally, the three structures show an integrated, sequential role in memory consolidation and retrieval (Izquierdo and Medina, 1993; Izquierdo et al., 1997) as well as coordinated activity and plasticity (Yaniv et al., 2003). For example, retrograde amnesia has been shown in rats after infusion of a GABA_A_ receptor agonist or a glutamate AMPA receptor antagonist, given immediately (but not at 30 min) post-training into the hippocampus and amygdala, and at 30-180 min (but not immediately) into the entorhinal cortex (Izquierdo et al., 1997).

The identified A-MTL subsystems could be further differentiated by their iFC peak. These peaks were located in distinct structures of the A-MTL along its longitudinal axis (Figure 2A), which were, to a large extent (i.e., for the majority of A-MTL subsystems), replicated in two additional, independent samples (Figures 3 and 4; Tables S4 and S5). We interpret this finding as evidence for a distinctive anterior-posterior organization of subsystems in the A-MTL, with the peak location acting as each subsystem’s functional ‘anchor.’ Crucially, the results based on this data-driven approach are in line with previous neuroimaging reports of distinctive functional organization among core MTL structures (Kahn et al., 2008; Libby et al., 2012; Maass et al., 2015) or amygdala sub-regions (Roy et al., 2009) in humans. Our results also expand to the entire A-MTL the previous data-driven demonstration of discrete functional sub-regions along the human hippocampus (Blessing et al., 2016). Furthermore, our longitudinally arranged A-MTL subsystems also match the longitudinally organized core MTL subsystems described in rodents and non-human primates as defined by other methods (for particular examples, see, e.g, Strange et al., 2014). Detailed comparative studies should, however, determine whether A-MTL subsystems obtained from functional connectivity in humans overlap with genetically or anatomically defined A-MTL subsystems in animal models.

#### 4.1.2 Global cortical and subcortical connectivity patterns mirror the local organization of the A-MTL

The twelve A-MTL subsystems we found are embedded into global-iFC patterns that follow, at a cortico-subcortical level, the longitudinal A-MTL organization (Figure 2B). Global-iFC patterns are located in medial insular-cingulate and basal ganglia/thalamus/hypothalamus regions that overlap with medial cortical and subcortical brain networks: the default mode, salience, basal ganglia/thalamus, and hypothalamus/brainstem networks. These cortico-subcortical brain networks, associated with distinct functional domains, have been recently suggested to collectively modulate allostatic-interoceptive functions in humans (Barrett and Simmons, 2015; Kleckner et al., 2017). Next, we will describe these A-MTL subsystems and their embedding in the global-iFC patterns in more detail.

### 4.2 Single A-MTL subsystems and their embedding into global-iFC patterns

#### 4.2.1 Anterior/frontoinsular A-MTL subsystems

The three anterior/frontoinsular A-MTL subsystems were centered on the amygdala, the anterior hippocampus, and the perirhinal cortex, and connected to the anterior and central hippocampus, and the entorhinal and parahippocampal cortices. At the global level, these subsystems included the anterior cingulate, orbitofrontal, and insular cortices; the putamen, ventral striatum, ventral pallidum, cerebellum, and periaqueductal gray; and the temporal pole. Note that the spatial resolution of our rs-fMRI data impeded a more reliable anatomical characterization at the level of the amygdala nuclei (see *Limitations*). Previous studies in humans have shown similar local and global patterns of connectivity for the perirhinal cortex (Kahn et al., 2008; Libby et al., 2012; Wang et al., 2016) and the amygdala (Roy et al., 2009), though separately. For example, the perirhinal cortex has been shown to exhibit preferential connectivity with the anterior hippocampus as well as with an anterior temporal and frontal cortical network (Libby et al., 2012). The amygdala, in turn, has shown organized connectivity between its basolateral division and the hippocampus, parahippocampal gyrus, and superior temporal gyrus; between its centromedial division and striatum, insula, cerebellum, and dorsal anterior cingulate cortex; and between its superficial division and the cingulate gyrus, hippocampus, caudate, and nucleus accumbens (Roy et al., 2009).

Previous structural connectivity evidence from animal studies (e.g., Van Hoesen and Pandya, 1975; Deacon et al., 1983; Carmichael and Price, 1995; McDonald et al., 1999; Stefanacci and Amaral, 2000) also supports our findings. For example, extensive reciprocal connections with the amygdala (e.g., medial and lateral, medial basal, accessory basal, and cortical nuclei) in the macaque have been shown for agranular and dysgranular insula (Mufson et al., 1981). Similar projections have also been shown for MTL regions such as the parahippocampal cortex, the superior temporal gyrus (Stefanacci and Amaral, 2000), the rostral entorhinal cortex, and – at a higher proportion – the rostral perirhinal cortex, which also receives back-projections from the amygdala (Deacon et al., 1983). Regions of the three anterior/frontoinsular A-MTL subsystems have previously been implied in semantically driven personal evaluations of social and asocial stimuli (Guo et al., 2013; Zhou and Seeley, 2014; Ranasinghe et al., 2016) as well as in emotional-autonomic responses, salience processing, and inhibitory control (Heimer and Van Hoesen, 2006; Seeley et al., 2007; Menon and Uddin, 2010), highly resembling a semantic-appraisal network and the aforementioned salience network, respectively.

#### 4.2.2 Anterior/subcortical A-MTL subsystems

Anterior/subcortical A-MTL subsystems were centered on anterior portions of the hippocampus and showed connectivity to the amygdala, central hippocampus, and entorhinal, perirhinal, and parahippocampal cortices. These subsystems were also functionally connected to ventromedial frontal and insular cortices, superior temporal gyrus, basal forebrain, nucleus accumbens, caudate nucleus, putamen, thalamus, hypothalamus, and upper pons. Previous studies in humans have also reported specific connectivity of the anterior hippocampus to the hypothalamus (Blessing et al., 2016); prefrontal cortex (Zarei et al., 2013); caudate, putamen, and nucleus accumbens (Qin et al., 2015); entorhinal and perirhinal cortices (Kahn et al., 2008); and amygdala and insula (Robinson et al., 2015).

Neuronal connectivity evidence from rodents and primates has indicated a set of descending projections from ventral hippocampal/subicular, amygdalar, and medial prefrontal cortical structures to the periventricular and medial zones of the hypothalamus involved in neuroendocrine, autonomic, and motivated behavior (Fanselow and Dong, 2010). Moreover, the connectivity of the anterior hippocampus to the ventral striatum and the mesolimbic dopamine system confers a role to the A-MTL in goal-directed behavior (Pennartz et al., 2011; Strange et al., 2014). Regions of these anterior/subcortical A-MTL subsystems are involved in reward-motivated behavior (Haber and Knutson, 2010), pain modulation (Zambreanu et al., 2005; Tracey and Mantyh, 2007), and complex motor and non-motor behavior (DeLong and Wichmann, 2007; Haber and Calzavara, 2009).

#### 4.2.3 Posterior/default mode network A-MTL subsystems

Posterior/default mode network A-MTL subsystems were anchored to portions of the anterior hippocampus, as well as the central hippocampus, and the posterior parahippocampal cortex. These subsystems connected to the amygdala, the whole extension of the hippocampus, and entorhinal, perirhinal, and parahippocampal cortices. Additionally, these subsystems were functionally connected to ventromedial, and dorsomedial frontal cortex, subgenual anterior and posterior cingulate cortices, middle temporal gyrus, retrosplenial cortex, inferior parietal lobule, precuneus, occipital cortex, nucleus accumbens, hypothalamus, cerebellum, and lower brain stem. This organization confirms previous reports in other studies in humans (Kahn et al., 2008; Libby et al., 2012; Qin et al., 2015; Wang et al., 2016). For example, the connectivity with the default mode network along the parahippocampal gyrus is characterized by being dominantly posterior (Qin et al., 2015). In other words, compared to its anterior portions (i.e., perirhinal cortex), the posterior portions of the parahippocampal gyrus (i.e., parahippocampal cortex) show stronger connectivity with default mode regions – such as the posterior cingulate cortex, retrosplenial cortex, or inferior temporal gyrus – (Wang et al., 2016). Middle and posterior parts of the hippocampus show a similar posterior-predominant pattern with posterior cingulate cortex (Zarei et al., 2013; Wang et al., 2016), whereas anterior parts of the hippocampus show this predominance with the prefrontal cortex (Zarei et al., 2013).

In humans, there is evidence of structural connectivity between the core MTL and the posterior component of the default mode network (i.e., the retrosplenial cortex) (Greicius et al., 2009). Similarly, the rostrocaudal topography of hippocampal projections compiled across primate studies indicates that projections to retrosplenial, anterior cingulate, or inferior temporal cortices arise, predominantly, from caudal (or posterior) portions of the hippocampus (Aggleton, 2012). Regions of the identified posterior/default mode networks and their interaction with the MTL have been associated, in general, with cognitive processes such as spatial navigation, planning, and semantic and episodic memory (Buckner et al., 2008).

### 4.3 Functional implications

Our findings indicate a functional organization for the whole MTL that goes beyond previous proposals by showing the critical involvement of the amygdala in all A-MTL subsystems. Moreover, the connectivity peaks found with our data-driven mICA approach highlight a longitudinal organization along the A-MTL that can further provide exact seed locations to inform future studies based on, e.g., structural MRI.

Beyond such local integration within the MTL, our results could complement – at large-scale systems level – the framework of a recently proposed ‘allostatic-interoceptive system’ (Barrett and Simmons, 2015; Kleckner et al. 2017). The allostatic-interoceptive system has been described as a domain-general brain system that includes visceromotor limbic – such as the cingulate cortices, ventral anterior insula, posterior orbitofrontal cortex, temporal pole, and parahippocampal gyrus – and primary interoceptive cortices – like the mid and posterior insula – as well as subcortical structures – like striatum and hypothalamus – (Barrett and Simmons, 2015; Chanes and Barrett, 2016). These regions correspond to exactly the same cortical and subcortical A-MTL subsystems described in the present study: the ‘anterior/frontoinsular,’ the ‘anterior/subcortical,’ and the ‘posterior/default mode network’ A-MTL subsystems. From a functional perspective, the allostatic-interoceptive system has been proposed to underpin functions that link the control of homeostatic body-oriented processes (i.e., interoception) with the body-referenced control of behavioral interactions of the organism with the environment (i.e., allostasis). In brief, the allostatic-interoceptive system matches the body’s physiology with its behavior. Such system, in turn, might constitute a unifying neural model integrating typical core-MTL and amygdala functions such as spatial navigation/memory consolidation and biological significance/emotional processing.

### 4.4 Limitations

In interpreting our results, several limitations must be taken into account. First, the spatial resolution used in our study prevented a solid outline of the sub-regional involvement within the A-MTL. However, the current study was a first attempt at characterizing subsystems of the whole A-MTL and determining whether these subsystems cover both amygdala and core MTL regions alike, showing good replicability across different samples. Second, it is unclear what type of connectivity is reflected by BOLD iFC. Therefore, our results share the same limitations derived from BOLD imaging. However, recent evidence shows that neuronal co-activations among subsets of cortical areas precede hemodynamic signal co-activations (i.e., the basis of BOLD iFC) among the same subsets of areas (Matsui et al., 2016). Future studies using intracranial electroencephalography in patients with MTL epilepsy could help to shed more light on this issue.

## 5 Conclusion

In summary, *in-vivo* data-driven intrinsic functional connectivity in humans revealed subsystems of the medial temporal lobe including the amygdala that could be reproduced in different samples of cognitively normal adults. These subsystems consistently covered parts of the amygdala, hippocampus, and entorhinal cortex, and showed a discrete longitudinal organization along the medial temporal lobes. This distinctive anterior-posterior organization was also found at the level of the whole brain, with local subsystems being functionally connected to frontoinsular, subcortical, or default mode networks – all of which have been described as part of an allostatic-interoceptive system.

## Conflict of Interest

The authors declare no competing conflict of interests.

## Acknowledgments

This work was supported by the European Union FP7 Marie Curie ITN Grant 606901 (INDIREA) and the German Academic Foundation. We thank Dr. Mihai Avram for providing the preprocessed data of the first replication sample.

