## Supplementary Material for "Human subsystems of medial temporal lobes extend locally to amygdala nuclei and globally to an allostatic-interoceptive system"

**Content:**

- *Separation of neural from non-neural local-iFC patterns*
  - Figure S1. Non-neural iFC patterns
  - Table S1. Percentage of voxels in CSF for the global-iFC patterns obtained from mICA with 20 dimensions
- *Control analyses: number of ICA dimensions and replication*
  - Number of ICA dimensions
    - Figure S2. Control analysis to determine the optimal ICA dimensionality
    - Table S2. Spatial cross-correlations of (global; local) iFC patterns derived from mICA
    - Table S3. Peaks of iFC patterns derived from mICA (10, 20, and 30 dimensions)
  - Replication
    - Supplementary Materials and Method
      - Replication sample 1
      - Replication sample 2
    - Supplementary Results
      - Table S4. Correspondence between the global and local iFC patterns of the original dataset and replication sample 1 (REP. 1)
      - Table S5. Correspondence between the global and local iFC patterns of the original dataset and replication sample 2 (REP. 2)
- *Summary of local-iFC pattern schematics of Figure 2B*
  - Figure S3. Percentage of overlap among all local-iFC patterns.
- *References*

***Separation of neural from non-neural local-iFC patterns***

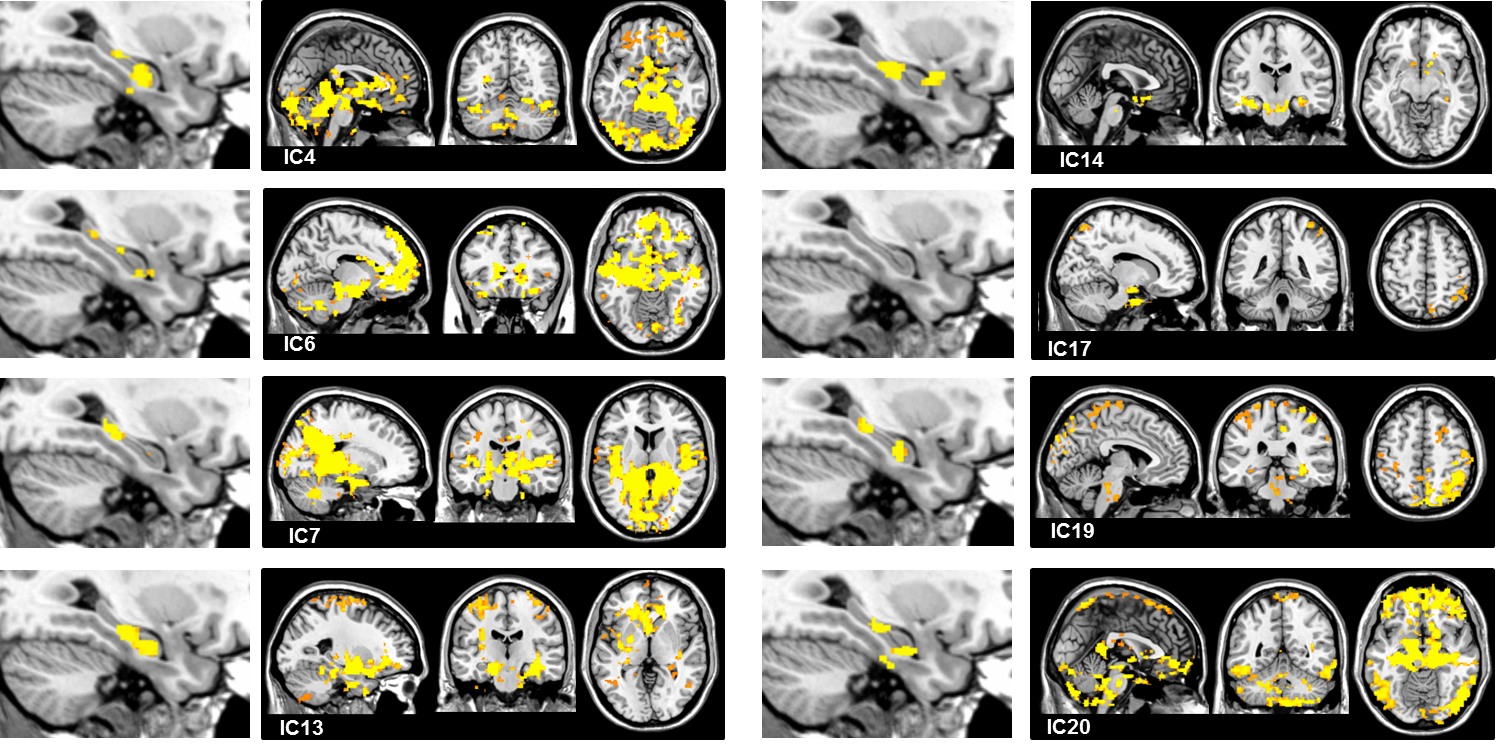

**Figure S1. Non-neural iFC patterns.** Local- and (corresponding) global-iFC patterns of the A-MTL identified as ‘non-neural’ (i.e., patterns with a percentage of voxels in CSF at or above the group mean [from all global-iFC patterns]). In yellow, significant voxels of global iFC (p-values; p < 0.05 FWE corrected for multiple comparisons).

**Table S1.** Percentage of voxels in CSF for the global-iFC patterns obtained from mICA with 20 dimensions

| **Global-iFC pattern (IC)** | **Total percentage of voxels in CSF** | **Label** |
| --- | --- | --- |
| IC1 | 5% | ‘Neural’ |
| IC2 | 12% | ‘Neural’ |
| IC3 | 4% | ‘Neural’ |
| IC4 | 29% | ‘Non-neural’ |
| IC5 | 13% | ‘Neural’ |
| IC6 | 18% | ‘Non-neural’ |
| IC7 | 15% | ‘Non-neural’ |
| IC8 | 11% | ‘Neural’ |
| IC9 | 3% | ‘Neural’ |
| IC10 | 5% | ‘Neural’ |
| IC11 | 8% | ‘Neural’ |
| IC12 | 13% | ‘Neural’ |
| IC13 | 20% | ‘Non-neural’ |
| IC14 | 27% | ‘Non-neural’ |
| IC15 | 13% | ‘Neural’ |
| IC16 | 10% | ‘Neural’ |
| IC17 | 20% | ‘Non-neural’ |
| IC18 | 9% | ‘Neural’ |
| IC19 | 34% | ‘Non-neural’ |
| IC20 | 14% | ‘Non-neural’ |

Mean ± standard deviation percentage of voxels in CSF = 14% ± 8%

***Control analyses: number of ICA dimensions and replication***

*Number of ICA dimensions*

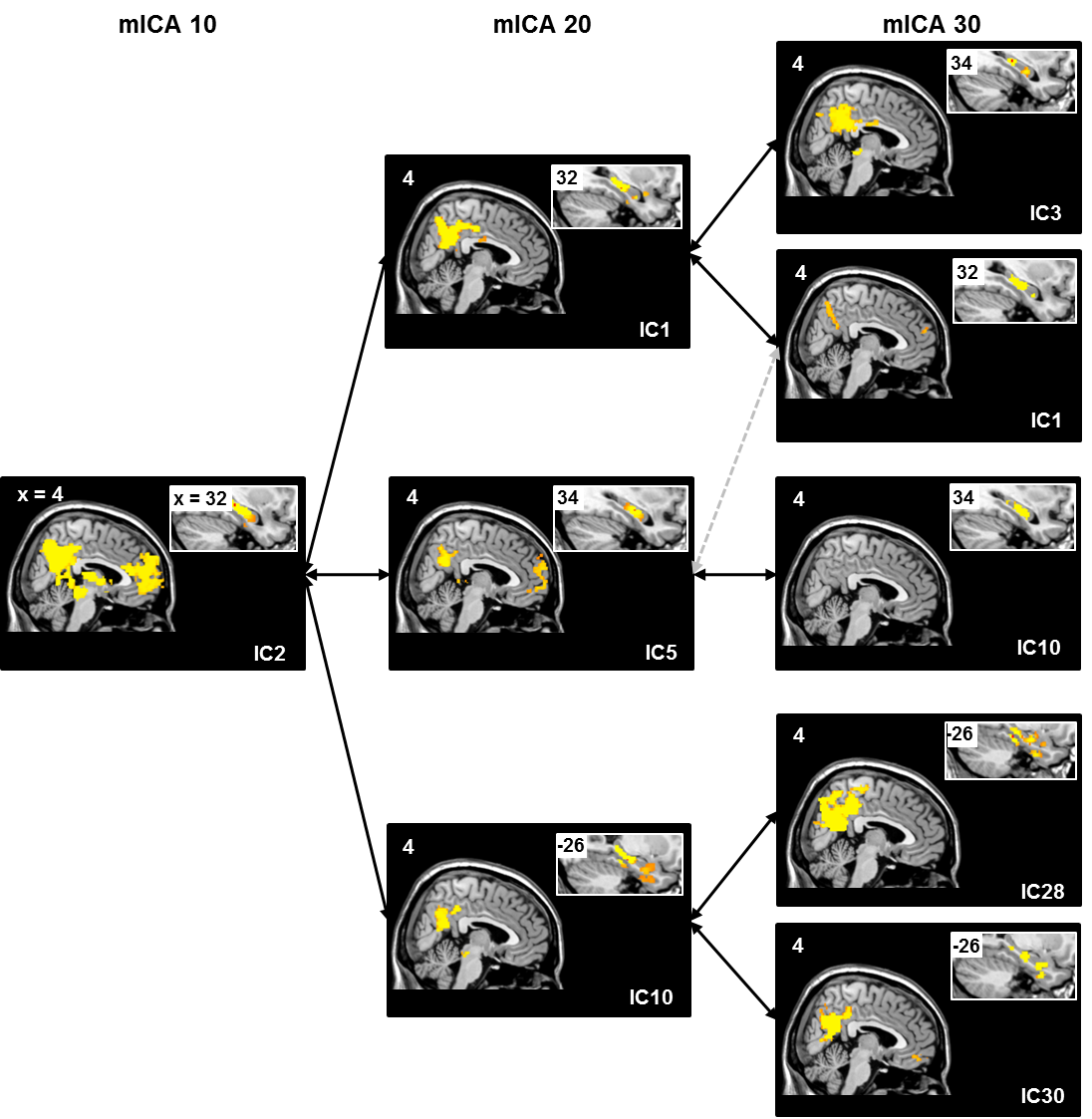

**Figure S2. Control analysis to determine the optimal ICA dimensionality.** Example of local- and global-iFC patterns identified as neural shown for three masked ICAs (mICA) with 10 (first column), 20 (second column), and 30 (third column) dimensions. In yellow, voxels significantly belonging to the iFC patterns (p-values; p < 0.05 FWE corrected for multiple comparisons). See Table S2 for the correlation coefficients and Table S3 for the peaks in each dimensionality.

**Table S2.** Spatial cross-correlations of (global; local) iFC patterns derived from mICA

| **10 ICs** | **20 ICs** | **30 ICs** |
| --- | --- | --- |
| **1.** IC2 | **1.1** IC1 (r = 0.46; r = 0.45)  **1.2** IC5 (r = 0.40; r = 0.56)  **1.3** IC10 (r = 0.38; r = 0.53) | **1.1.1** IC3 (r = 0.47; r = 0.38)  **1.1.2.** IC1 (r = 0.43; r = 0.74)  **1.2.3** IC10 (r = 0.35; r = 0.51)  * IC1 (r = 0.13; r = 0.36)  **1.3.4** IC28 (r = 0.39; r = 0.35)  **1.3.5** IC30 (r = 0.28; r = 0.31) |
| **2.** IC4 | **2.4** IC8 (r = 0.53; r = 0.40)  **2.5** IC12 (r = 0.40; r = 0.28)  **2.6** IC18 (r = 0.38; r = 0.44) | * IC28 (r = 0.49; r = 0.40)  * IC30 (r = 0.42; r = 0.52)  * IC30 (r = 0.37; r = 0.44)  * IC28 (r = 0.25; r = 0.50)  * IC28 (r = 0.53; r = 0.39)  * IC30 (r = 0.38; r = 0.43) |
| **3.** IC5 | * IC5 (r = 0.37; r = 0.52)  * IC12 (r = 0.42; r = 0.46)  **3.7** IC9 (r = 0.31; r = 0.46) | --  --  --  --  **3.7.6** IC7 (r = 0.45; r = 0.59)  **3.7.7** IC25 (r = 0.34; r = 0.40) |
| **4.** IC6 | **4.8** IC15 (r = 0.56; r = 0.54)  **4.9** IC11 (r = 0.37; r = 0.39)  * IC9 (r = 0.33; r = 0.59) | **4.8.8** IC19 (r = 0.40; r = 0.44)  **4.8.9** IC6 (r = 0.28; r = 0.37)  **4.9.10** IC9 (r = 0.66; r = 0.56)  * IC25 (r = 0.49; r = 0.40)  --  -- |
| **5.** IC7 | **5.10** IC3 (r = 0.45; r = 0.59)  * IC15 (r = 0.34; r = 0.52)  **5.11** IC2 (r = 0.31; r = 0.57) | **5.10.11** IC5 (r = 0.36; r = 0.50)  * IC10 (r = 0.30; r = 0.48)  --  --  * IC6 (r = 0.62; r = 0.72)  * IC19 (r = 0.41; r = 0.51) |

* ‘Repeated’ IC, i.e., that correlates with more than one of the ICs in the previous column. The coefficients in the second column correspond to the correlation with the ICs in the first column; those in the third column correspond to the correlations with the ICs in the second column. The correlation coefficient of the global iFC is shown first. IC: Independent component; ICA: independent component analysis. The first number in the numbering sequence reflects the sequence of mICA 10, the second that of mICA 20, and the third that of mICA 30. Repeated ICs are not included in the numbering sequence within a particular mICA. Only spatial cross-correlations among neural iFC patterns are shown.

**Table S3.** Peaks of iFC patterns derived from mICA (10, 20, and 30 dimensions)

| **10 ICs** | **20 ICs** | **30 ICs** |
| --- | --- | --- |
| **1.** IC2: 32, -30, -8 | **1.1** IC1: 32, -26, -14  **1.2** IC5: 34, -20, -14  **1.3** IC10: -26, -28, -14 | **1.1.1** IC3: 34, -30, -8 / **1.1.2** IC1: 32, -26, -14  **1.2.3** IC10: 36, -18, -16 / * IC1: 32, -26, -14  **1.3.4** IC28: -26, -34, -16 / **1.3.5** IC30: 28, -28, -20 |
| **2.** IC4: 28, -26, -14 | **2.4** IC8: 20, -32, -14  **2.5** IC12: -20, -16, -18  **2.6** IC18: -24, -32, -18 | * IC28: -26, -34, -16 / * IC30: 28, -28, -20  * IC30: 28, -28, -20 / * IC28: -26, -34, -16  * IC28: -26, -34, -16 / * IC30: 28, -28, -20 |
| **3.** IC5: -26, -22, -16 | * IC5: 34, -20, -14  * IC12: -20, -16, -18  **3.7** IC9: -26, -14, -18 | --  --  **3.7.6** IC7: -26, -12, -20 / **3.7.7** IC25: -24, -10, -36 |
| **4.** IC6: -24, -4, -16 | **4.8** IC15: -26, -4, -16  **4.9** IC11: -24, -8, -36  * IC9: -26, -14, -18 | **4.8.8.** IC19: 26, -8, -16 / **4.8.9.** IC6: -24, -14, -16  **4.9.10.** IC9: -24, -6, -34 / * IC25: -24, -10, -36  -- |
| **5.** IC7: 28, -12, -20 | **5.10** IC3: 30, -12, -20  * IC15: -26, -4, -16  **5.11** IC2: 22, -12, -18 | **5.10.11.** IC5: 30, -12, -20 / * IC10: 36, -18, -16  --  * IC6: -24, -14, -16 / * IC19: 26, -8, -16 |

* ‘Repeated’ IC, i.e., that correlates with more than one of the ICs in the previous column as in Table S2. The first number in the numbering sequence reflects the sequence of mICA 10, the second that of mICA 20, and the third that of mICA 30. Repeated ICs are not included in the numbering sequence within a particular mICA. IC: Independent component; ICA: independent component analysis.

*Replication*

*Supplementary Materials and Method*

Replication sample 1

Imaging data from sixty-one healthy adults from 18 to 65 years old (mean age: 36.4 ± 13.6 years; 18 females) were obtained from the Center for Biomedical Research Excellence (COBRE; <http://fcon_1000.projects.nitrc.org/indi/retro/cobre.html>) and were used as the first replication sample (REP. 1). Exclusion criteria were DSM-IV diagnoses of any kind, psychiatric or neurological disorders, history of substance abuse, and significant head trauma (also see <http://fcon_1000.projects.nitrc.org/indi/retro/cobre.html>). From the original sample of 74 participants, 2 had to be excluded due to history of mental disorders and 11 were excluded due to significant head motion. For intrinsic functional connectivity (iFC) analysis, 150 volumes were collected on a 3T Siemens Trio scanner during subjects’ rest, with a 12-channel radio frequency coil, using single-shot full k-space echo-planar imaging (EPI) with ramp sampling correction and the intercomissural line (AC-PC) as a reference; Repetition time, TR = 2,000 ms; Time to Echo, TE = 29 ms; matrix size = 64 x 64, 32 slices; voxel size = 3.75 x 3.75 x 4.55 mm^3^. For co-registration of the rs-fMRI volumes, a high-resolution T1-weighted volume was also acquired using a 3D magnetization prepared rapid acquisition gradient echo (MPRAGE) sequence (TR = 2,530 ms; TEs = 1.64, 3.5, 5.36, 7.22, 9.08; TI, inversion time = 900 ms; flip angle: 7°; matrix size = 256 x 256, 176 slices; voxel size = 1 mm isotropic). Imaging data preprocessing and analysis were conducted as described in the main text.

Masked ICA within the A-MTL was performed with 20 components, which were the input for two dual regressions, one within the A-MTL (local-iFC patterns) and one in the whole brain (global-iFC patterns). Here we followed the same procedure to select local-iFC subsystems based on the global-iFC patterns that we described in the main text.

Replication sample 2

Imaging data from twenty-nine healthy young adults from 18 to 35 years old (mean age: 26.0 ± 4.1 years; 14 females) were obtained from an ongoing independent project on aging and visual attention, whose data were part of Ruiz-Rizzo et al. (2019), and were used as the second replication sample (REP. 2). Exclusion criteria were the same as those of the replication sample 1. For iFC analysis, 600 volumes were acquired on a Philips Ingenia 3T system (Netherlands) during subjects’ rest, using a 32-channel SENSE head coil using a multiband EPI sequence, with a 2-fold in-plane SENSE acceleration (SENSE factor, S = 2) and an M-factor of 2; TR = 1,250 ms; TE = 30 ms; matrix size = 64 x 64, 40 slices; voxel size = 3 x 3 x 3.29 mm^3^. For co-registration of the rs-fMRI volumes, a high-resolution T1-weighted MPRAGE volume was also acquired (TR = 9 ms; TE = 4 ms; TI = 0 ms; flip angle = 8º; matrix size = 240 x 240, 170 slices; voxel size = 1 mm isotropic). Imaging data preprocessing and analysis were conducted as described in the main text and in *Replication sample 1*.

*Supplementary Results*

Figure 3 of the main text shows the comparison of the *global*-iFC patterns for the three samples: the original reported in the main manuscript, the replication sample 1 (REP. 1) and the replication sample 2 (REP. 2). Table S4 and S5 list the spatial cross-correlation coefficients between the original global iFC-patterns and those of the replication sample 1 (S4) and the replication sample 2 (S5). Note that for both replication samples, the global-iFC patterns were selected if their percentage of voxels in CSF was below the respective group mean (i.e., as in the original sample). In general, all global-iFC patterns were replicated in both samples (REP. 1: first and second column of Table S4; REP. 2: first and second column of Table S5). The ‘IC18-Dorsal parietal’ and the ‘IC2-Thal+BG’ of the original sample were the two global-iFC patterns with the highest correlation coefficients in both replication samples, whereas the ‘IC9-Left ventral’ was among the global patterns with the lowest correlation coefficients in both replication samples. As for the impact of sample size, the cross-correlation coefficients were relatively similar between replication samples (i.e., means: 0.30 and 0.27 for REP. 1 and REP. 2, respectively).

Figure 4 of the main text includes the *local*-iFC patterns of the original and both replication samples (see Tables S4 and S5 for the respective correlation coefficients). Overall, the spatial correspondence was better for the local-iFC than for the global-iFC patterns. The two local-iFC patterns with the highest correlation coefficients were ‘IC18-Dorsal parietal’ for both samples (as for the global-iFC patterns), and ‘IC2-Thal+BG’ and ‘IC1-Parietal right’ for REP. 1 and ‘IC3-Brain stem’ for REP. 2. Like with the global-iFC patterns, the local-iFC pattern corresponding to ‘IC9-Left ventral’ had the lowest cross-correlation coefficient in REP. 2, whereas ‘IC10-Parietal left’ had it in REP. 1 and ‘IC11-ACC’ had it in both samples. This time, the replication sample with a higher size (REP. 1) had, on average, slightly higher cross-correlation coefficients than the smaller one (REP. 1: 0.49 vs. REP. 2: 0.44). Finally, the iFC peaks of the replication samples’ A-MTL subsystems were also remarkably close to the iFC peaks described for the original sample (Table S3), as can be observed in the two last columns of Tables S4 and S5.

**Table S4.** Correspondence between the global and local iFC patterns of the original dataset and replication sample 1 (REP. 1)

| **Original sample** | **Highest correlation coefficient** | **iFC peak in original sample** | **iFC peak in REP. 1** |
| --- | --- | --- | --- |
| IC15-Frontoinsular | [IC2]  r = 0.39 (g)  r = 0.50 (l) | -26, **-4**, -16 | 26, **-4**, -20 |
| IC16-Orbitofrontal | [IC14]  r = 0.23  r = 0.48 | 26, **-6**, -24 | -22, **-32**, -20 |
| IC11-ACC | [IC11]  r = 0.24  r = 0.37 | -24, **-8**, -36 | -24, **-14**, -32 |
| IC3-Brain stem | [IC8]  r = 0.24  r = 0.56 | 30, **-12**, -20 | 26, **-8**, -18 |
| IC2-Thal+BG | [IC2]*  r = 0.44  r = 0.58 | 22, **-12**, -18 | 26, **-4**, -20 |
| IC9-Left ventral | [IC11]*  r = 0.18  r = 0.41 | -26, **-14**, -18 | -24, **-14**, -32 |
| IC12-Ventral frontal and parietal | [IC15]  r = 0.39  r = 0.52 | -20, **-16**, -18 | 24, **-28**, -20 |
| IC5-Dorsal frontal and parietal | [IC15]*  r = 0.20  r = 0.44 | 34, **-20**, -14 | 24, **-28**, -20 |
| IC1-Parietal right | [IC9]  r = 0.25  r = 0.59 | 32, **-26**, -14 | -34, **-26**, -10 |
| IC10-Parietal left | [IC14]*  r = 0.31  r = 0.39 | -26, **-28**, -14 | -22, **-32**, -20 |
| IC8-Ventral parietal | [IC14]*  r = 0.33  r = 0.49 | 20, **-32**, -14 | -22, **-32**, -20 |
| IC18-Dorsal parietal | [IC14]*  r = 0.44  r = 0.60 | -24, **-32**, -18 | -22, **-32**, -20 |

* ‘Repeated’ IC: i.e., that correlates with more than one of the ICs in the previous column; G: global iFC; l: local iFC. Peaks in x, y, z MNI coordinates (in mm). See corresponding figures in the main text (Figure 3 for global-iFC patterns and Figure 4 for local-iFC patterns).

**Table S5.** Correspondence between the global and local iFC patterns of the original dataset and replication sample 2 (REP. 2)

| **Original sample** | **Highest correlation coefficient** | **iFC peak in original sample** | **iFC peak in REP. 2** |
| --- | --- | --- | --- |
| IC15-Frontoinsular | [IC2]  r = 0.33 (g)  r = 0.41 (l) | -26, **-4**, -16 | 24, **-6**, -16 |
| IC16-Orbitofrontal | [IC10]  r = 0.26  r = 0.53 | 26, **-6**, -24 | 22, **-10**, -14 |
| IC11-ACC | [IC3]  r = 0.16  r = 0.28 | -24, **-8**, -36 | 28, **-10**, -20 |
| IC3-Brain stem | [IC3]*  r = 0.22  r = 0.68 | 30, **-12**, -20 | 28, **-10**, -20 |
| IC2-Thal+BG | [IC2]*  r = 0.41  r = 0.52 | 22, **-12**, -18 | 24, **-6**, -16 |
| IC9-Left ventral | [IC3]*  r = 0.19  r = 0.21 | -26, **-14**, -18 | 28, **-10**, -20 |
| IC12-Ventral frontal and parietal | [IC12]  r = 0.20  r = 0.46 | -20, **-16**, -18 | -26, **-30**, -20 |
| IC5-Dorsal frontal and parietal | [IC3]*  r = 0.22  r = 0.40 | 34, **-20**, -14 | 28, **-10**, -20 |
| IC1-Parietal right | [IC12]*  r = 0.17  r = 0.35 | 32, **-26**, -14 | -26, **-30**, -20 |
| IC10-Parietal left | [IC12]*  r = 0.28  r = 0.34 | -26, **-28**, -14 | -26, **-30**, -20 |
| IC8-Ventral parietal | [IC12]*  r = 0.35  r = 0.51 | 20, **-32**, -14 | -26, **-30**, -20 |
| IC18-Dorsal parietal | [IC12]*  r = 0.43  r = 0.61 | -24, **-32**, -18 | -26, **-30**, -20 |

* ‘Repeated’ IC: i.e., that correlates with more than one of the ICs in the previous column. G: global iFC; l: local iFC. Peaks in x, y, z MNI coordinates (in mm). See corresponding figures in the main text (Figure 3 for global-iFC patterns and Figure 4 for local-iFC patterns).

***Summary of local-iFC pattern schematics of Figure 2B***

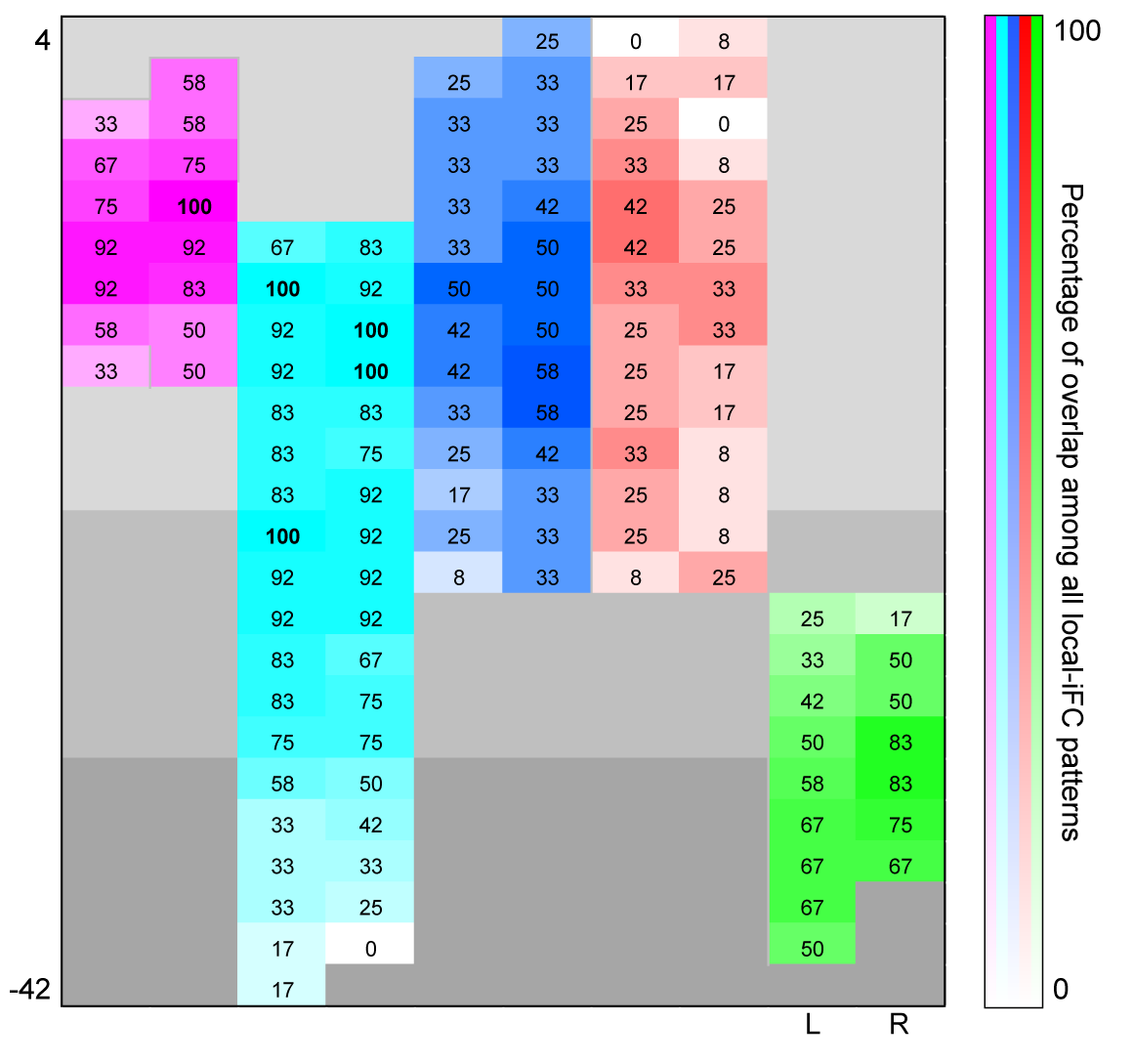

**Figure S3.** **Percentage of overlap among all local-iFC patterns.** This figure summarizes the local-iFC pattern schematics of Figure 2B (main text) by showing the slice-wise percentage of overlap (for each A-MTL structure and each hemisphere) among all local-iFC patterns. Note that the color coding corresponds to the structure labels on the top left of Figure 2A (main text).
